# Visual boundary cues, alone, can anchor place and grid cells in virtual reality

**DOI:** 10.1101/2023.10.27.564224

**Authors:** Xiuting Yang, Francesca Cacucci, Neil Burgess, Thomas Wills, Guifen Chen

## Abstract

The hippocampal formation contains neurons responsive to an animal’s current location and orientation, which together provide the organism with a neural map of space^1–3^. Spatially tuned neurons rely on external landmark cues and internally generated movement information to estimate position^4,5^. An important class of landmark cue are the boundaries delimiting an environment, which can define place cell field position^6,7^ and stabilise grid cell firing^8^. However, the precise nature of the sensory information used by the hippocampus to detect boundaries remains unknown. Here we use 2-dimensional virtual reality (VR)^9^ to show that purely visual cues from the surrounding walls are both sufficient and necessary to stabilise place and grid cell responses. Unstable grid cells retain internally coherent, hexagonally arranged firing fields, but these fields ‘drift’ with respect to the virtual environment over periods >5 sec. Optic flow from a virtual floor does not slow drift dynamics, emphasising the importance of boundary-related visual information. Surprisingly, place fields are more stable close to boundaries even when these are invisible. Invisible boundary locations can only be inferred from the relative movements of the animal and a reward beacon, specifically, from the mismatch between observed beacon movement and that predicted from motor output. This shows that boundary information can be detected through sensorimotor feedback. We find that subsets of place cells show allocentric directional tuning towards the beacon, with strength of tuning correlating with place field stability when boundaries are removed.

## RESULTS

Place and grid cells (796 place cells and 138 grid cells) were recorded while mice navigated in a 2-dimensional (2D) virtual reality environment^9^ (Figure S1A&B): the external sensory cues defining the environment were purely visual, while the animal’s movement around the environment was coupled to its movement on a spherical trackball. Mice were trained to navigate towards a moveable beacon to receive a reward^9^ (see methods). After mice had learned this task^9^, we tested the necessity of different visual cues for place and grid cell firing, by removing specific subsets of these cues as mice continued to perform the navigation task (‘Cue-absent’ trials). In three different manipulations (Figure S1), different subsets of cues were removed: a) the walls of the virtual environment (‘Walls Off’), b) the patterned floor of the environment (‘Floor Off’), c) both the walls and the floor (‘All Off’). In the cue-absent trials, the removed cues were replaced by a plain dark grey background. In all manipulations, the beacon indicating the current reward position remained visible and moved to a pseudo-random location after the mouse had consumed the reward (see Methods for more details). For each manipulation, a baseline trial was carried out where all visual cues were present (a ‘cue-present’ trial, see Figure S1).

Removing the wall cues alone produced a strong disruption in the spatial specificity and within-trial stability of place cells, as did removing both walls and floor cues simultaneously. By contrast, removing only floor cues had no significant effect on place cell spatial firing (Figure 1A, 1D, 1B, 1E; Spatial information: ANOVA Cue presence*Manipulation, F(2,617) = 26.94, p < 0.001, Simple Main Effects [SME] cue present vs absent, Walls Off, p < 0.001, Floor Off, p = 0.305, All Off, p < 0.001; Intra-trial stability: ANOVA Cue presence*Manipulation, F(2,617) = 90.36, p < 0.001, SME absent vs present, Walls Off p < 0.001, Floor Off, p = 0.091, All Off, p < 0.001). Removing the floor cues only did, however, cause a modest degree of remapping, as measured by the across-trial stability of place cell firing between the cue-present and cue-absent trials (Figure 1C, 1F; ANOVA Cue presence*Manipulation, F(2,617) = 114.60, p < 0.001, SME absent vs present, Walls Off, p < 0.001, Floor Off, p < 0.001, All Off, p < 0.001). This indicates a degree of saliency of floor cues to place cell firing, but only the removal of wall cues resulted in a significant decrease in place cell stability.

**Figure 1.**
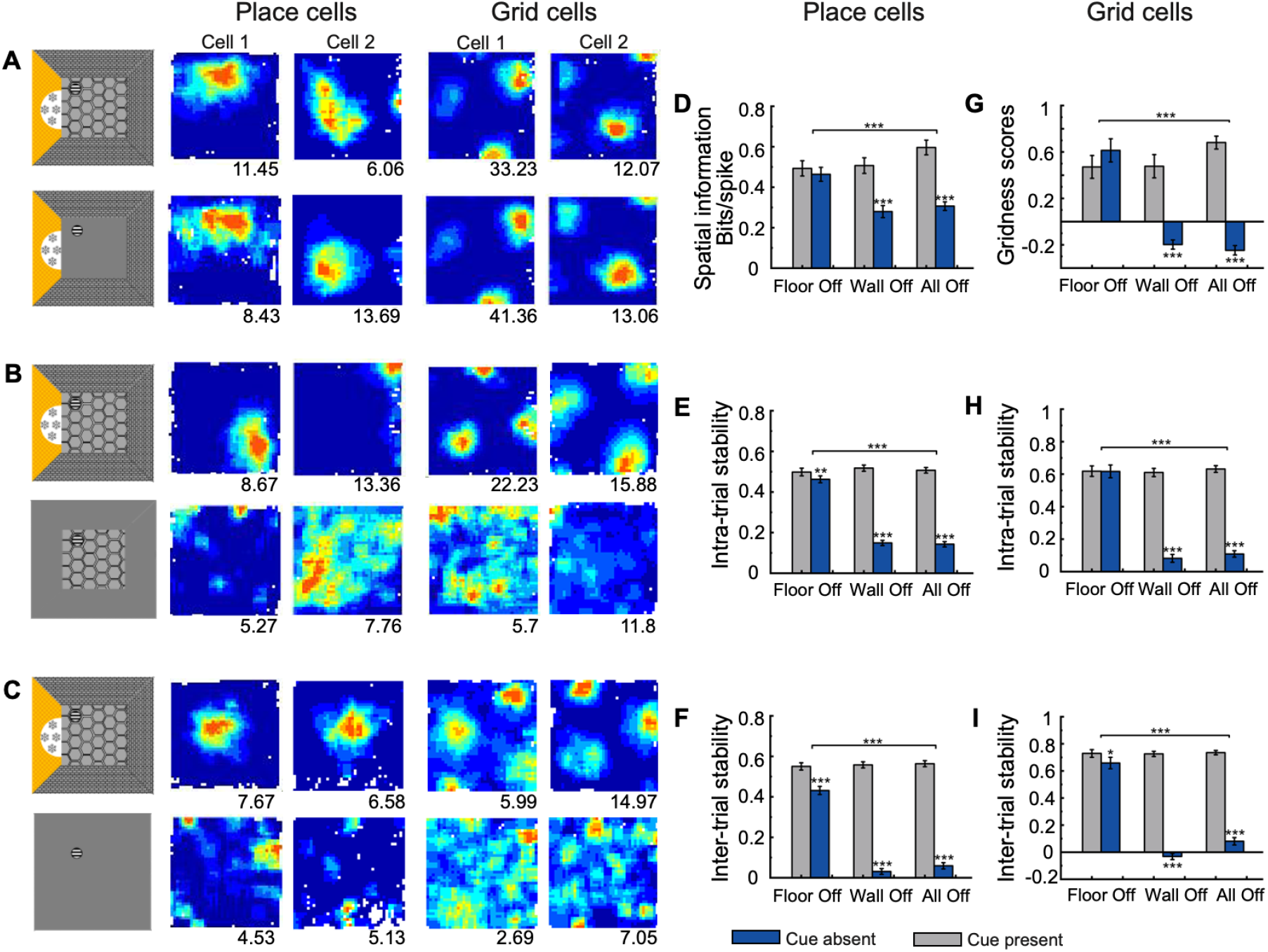
Visual cues from boundaries provide sufficient inputs to stabilise spatial patterns of place cells and grid cells. (A) Visual cues were removed from the floor (Floor Off). (B) Visual cues were removed from all four walls (Walls Off). (C) All environmental visual cues were removed (All Off). The panels on the left column are schematic of the manipulations with a cue-present environment on the top and a cue-absent environment on the bottom; the 2nd and 3rd columns are the rate maps of two example place cells in the corresponding environments; the 4th and 5th columns are rate maps of two example grid cells. Numbers at the bottom right of rate maps are peak firing rates (Hz). (D-I) Comparison of firing properties of place cells and grid cells in all three manipulations including spatial information (D), intra-trial stability (E) and inter-trial stability between the cue-present and cue-absent trials (F) for place cells; gridness scores (G), intra-trial stability (H) and inter-trial stability (I) for grid cells. ^**^ p < 0.01; ^***^ p < 0.001.

The effects of cue removal on grid cells paralleled those observed for place cells: removing walls only or floor and walls together resulted in a strong reduction in gridness scores and within-trial stability, while there was no significant effect of removing the floor cues alone (Figure 1A-1C, 1G-1I; Gridness score: ANOVA Cue presence*Manipulation, F(2,109) = 32.92, p < 0.001; SME absent vs present, Walls Off, p < 0.001, Floor Off, p = 0.202, All Off, p < 0.001; Intra-trial stability: ANOVA Cue presence*Manipulation, F(2,109) = 64.45, p < 0.001; SME absent vs present, Walls, p < 0.001, Floor, p = 0.972, All, p < 0.001). Notably, unlike place cells, grid cells did not significantly remap between cue-present and Floor-Off trials (ANOVA Cue presence*Manipulation, F(2,109) = 99.92, p < 0.001; SME absent vs present, Walls Off, p < 0.001, Floor Off, p = 0.079, All Off, p < 0.001).

Furthermore, the Walls-Off condition disrupted place and grid cell firing to the same extent as the All-Off condition: there were no significant differences in spatial information, gridness or intra-trial stability between these conditions (SME pairwise comparison: spatial information, p = 0.469; place cell intra-trial stability: p = 0.738; gridness, p = 0.487; grid cell intra-trial stability, p = 0.425). These results suggest that environmental walls are the cues responsible for stabilising place and grid cell firing within this virtual environment.

A subset of cells were also recorded in a second cue-present trial, directly after the cue-absent trial. Both place and grid cells restored their firing patterns after visual cues were replaced (Figure S2, see figure legend for detailed statistical results), indicating recording stability. As the Walls-Off and All-Off manipulations involved removing the visual cues from the proximal walls alongside the distal extra-maze cue (Figure S1), the loss of a stable directional heading in this condition could be a potential explanation for the observed reduction of place and grid cell stability. However, the directional tuning of co-recorded mEC head direction cells was not significantly reduced in either Walls-Off or All-Off conditions, suggesting consistent heading information was retained in these ‘Cue-absent’ conditions, even in the absence of the extra-maze cue (Figure S3).

Computational models predict that grid cells can generate regular hexagonal patterns from purely idiothetic information^10–13^. Anchoring these patterns to the external world using landmark cues is thought to prevent instability arising from accrued path integration error^14^. We tested whether, despite the breakdown of stable grid patterns when data were averaged over the whole trial (20 mins), grid patterns were present at short time scales in the cue-absent conditions, as this would be an indication of path integration drift. We focused on the Walls-Off and All-Off cue manipulations as these resulted in strong disruption of grid cell firing. Time-windowed auto-correlogram firing rate maps (see Methods) showed hexagonal patterns at short time intervals (Figure 2A, B; peaking at 5 secs), with progressive disruption at longer durations. Gridness scores of the time-windowed rate maps increased at longer durations when cues were present but decreased at longer durations when cues were absent (Figure 2C; 2-way ANOVA Duration-Cue presence: SME Duration F(5, 136) = 37.34, p < 0.001, interaction Duration*Cue presence F(5,136) = 13.87, p < 0.001). Thus, hexagonal grid firing patterns were detectable over short time intervals in both cue-absent conditions.

**Figure 2.**
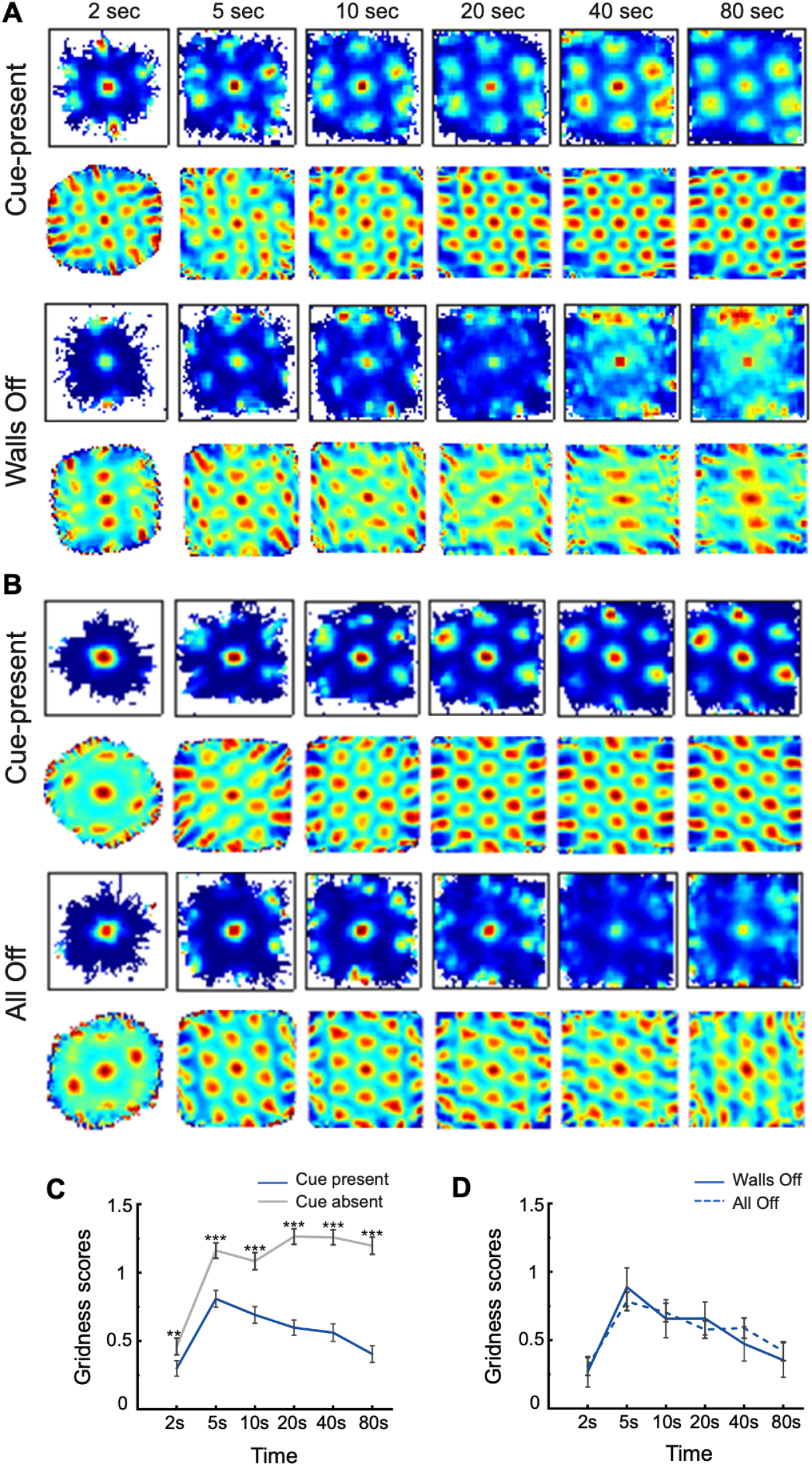
Grid cells maintain hexagonal patterns on short-time scales. (A-B) Time-windowed rate maps and their auto-correlograms with 2, 5, 10, 20, 40, and 80 s windows of representative grid cells in cue-present (top rows) and cue-absent (bottom rows) for the Wall-Off (A) and All-Off (B) conditions. (C) Mean (±SEM) elliptical gridness scores calculated from the time-windowed displacement firing rate maps comparing cue-present (gray) with cue-absent (blue) trials. (D) Mean (±SEM) elliptical gridness scores calculated from the time-windowed displacement firing rate maps comparing the Walls-Off (solid) with the All-Off (dotted) conditions. ^*^ p < 0.05; ^**^ p < 0.01; ^***^ p < 0.001.

Optic flow is thought to be an important source of movement information which can contribute to path integration^15,16^. However, the patterned visual information provided by the floor did not significantly improve the gridness score at short time scales (Figure 2D; Two-way ANOVA Manipulation-Duration: main effect of Manipulation (Walls Off vs All Off), F(5, 135) = 0.02, p = 0.904). There was no evidence, therefore, that optic flow from patterned visual information supports grid cell stability in the virtual reality environment.

In all cue-absent conditions, the animal’s movement was constrained within the same virtual boundaries as the cue-present trials, and the reward beacon remained visible at all times, with its position changing to pseudo-random locations after the animal consumed the reward. On average, the reward beacon moved approximately every 4-5 seconds, but was stable between these relocations. As the reward beacon was the only source of positional information in the All-Off condition, we reasoned that the mouse may infer the presence of the virtual boundary from the motion of the reward beacon relative to its own motor output. We tested whether such inferred boundaries could contribute to the short-time scale stability of spatial patterns we observed in the All-Off condition, by parsing the environment into boundary and centre zones, and calculating the intra-trial stability in each zone. We found that cue removal consistently reduced stability in the centre zone, relative to the boundary zone, for both place cells (Figure 3C; ANOVA Zone*Cue presence, F(1,611) = 6.97, p < 0.01) and grid cells (Figure 3D; ANOVA, Zone*Cue presence, F(1,106) = 34.97, p < 0.001). However, the specific effects of cue removal were different across the two cell types. Place cells were equally stable across environmental boundary and centre when cues were present, but were significantly more stable near boundaries in the All-Off condition (Figure 3C; Paired t-test: All Off, t(611) = 6.03, p < 0.001; cue-present, t(613) = 1.96, p = 0.051). These results suggest that, in a highly familiar environment, place fields can be at least partially stabilised by boundary cues even when these cues themselves are not perceptually available, but are rather inferred from the movement (relative to motor output) of other, perceptually available, cues. Grid cells, by contrast, were more stable in the environment centre when cues were present, but became equally unstable at all environmental locations in the All-Off condition (Figure 3D; Paired t-test: All Off, t(106) = 0.85, p = 0.40; Cue-present, t(106) = -8.41, p < 0.001). Taken together, this indicates that place, but not grid representations, might benefit from the presence of the moving reward beacons to calculate location.

**Figure 3.**
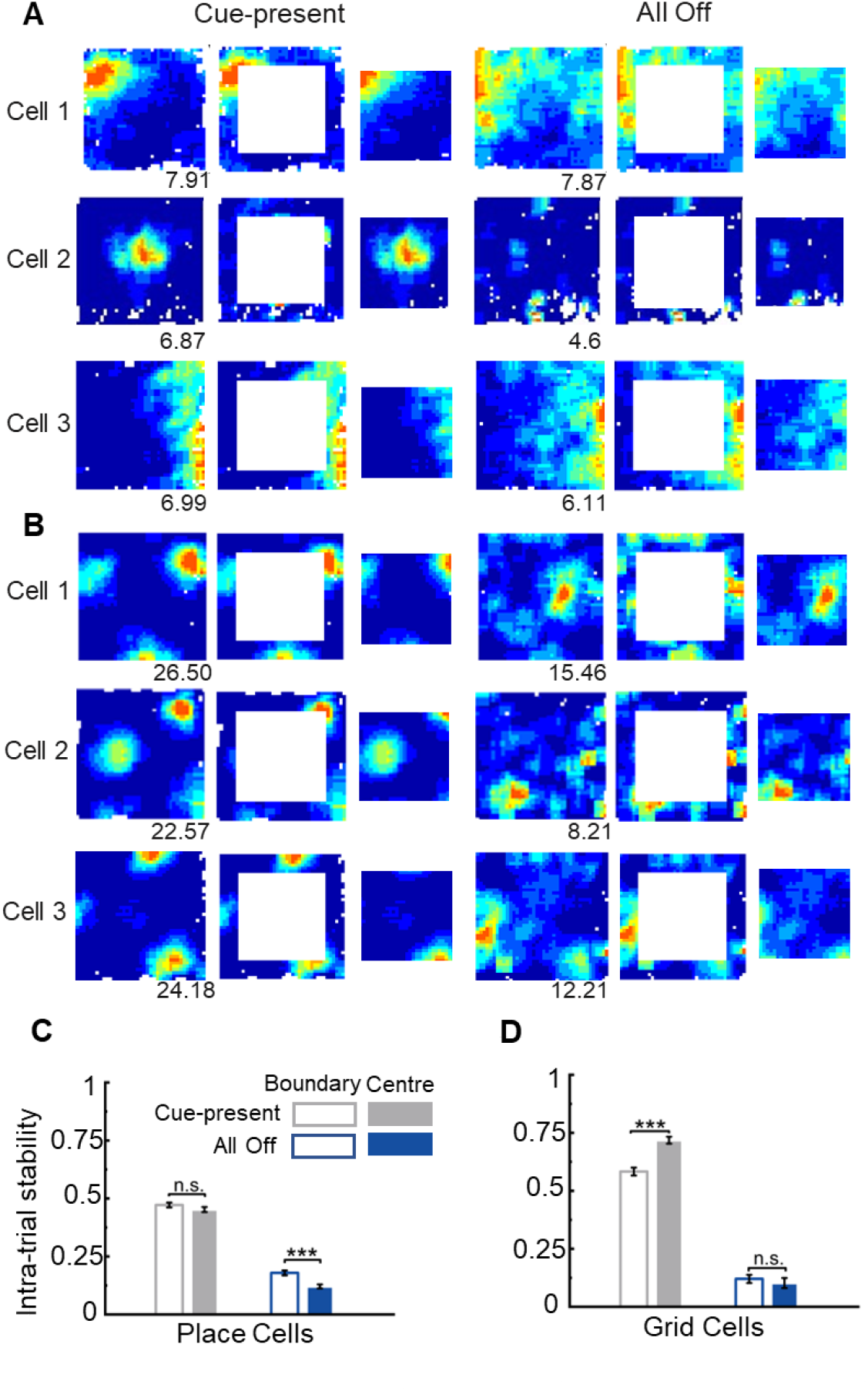
Proximity to inferred boundaries increases stability for both place cells and grid cells. (A) Rate maps of three example place cells recorded in a 60 cm virtual square in the cue-present and cue-absent (All-Off) conditions. (B) Rate maps of three example grid cells recorded in a 60 cm virtual square in the cue-present and All-Off conditions. The left column shows the rate maps for the whole environment, the middle column shows the boundary area of the maps, and the right column shows the central area of the maps. Numbers at the bottom right of firing rate maps are peak firing rate (Hz). (C) Intra-trial stability of boundary (open bars) and central (solid bars) rate maps in the cue-present (grey) and All-Off (blue) conditions for place cells. (D) Intra-trial stability of boundary (open bars) and central (solid bars) rate maps in the cue-present (grey) and All-Off (blue) conditions for grid cells. ^***^ p < 0.001.

Following our unexpected finding that place fields can be stabilised by a boundary inferred from the movement of a discrete cue (the reward beacon), we tested whether place cell firing may be modulated by this cue. Specifically, we tested whether place cells encoded the animal’s bearing to the reward beacon, irrespective of the animal’s head orientation. We found that, in the baseline (cue present) condition, 18.6% (114/614) of place cells showed significant directional tuning towards the reward beacons (see Methods) (Figure 4A-B). Directional tuning towards the beacon was reduced in the All-Off condition (Figure 4C; Cue-present: median = 0.12, IQR = 0.15; All-Off: median = 0.10, IQR = 0.12 for allocentric RV scores, Wilcoxon rank sum test, z = 7.30, p < 0.001), but there remained a correlation between the degree of beacon tuning in baseline and All-Off trials (Figure 4D; r = 0.44; p < 0.001, controlling for the boundary index) suggesting that beacon tuning is a cell-specific property. Moreover, the strength of place cells’ allocentric tuning (RV scores) in the All-Off condition was significantly and positively correlated with the intra-trial stability in the All-Off condition, irrespective of the relative position of place fields with respect to environmental boundaries (Figure 4E; partial correlation r = 0.19; p < 0.001, controlling for the boundary index), indicating that, in our experimental paradigm, the reward beacons help to stabilise the firing patterns of place cells in the All-Off condition.

**Figure 4.**
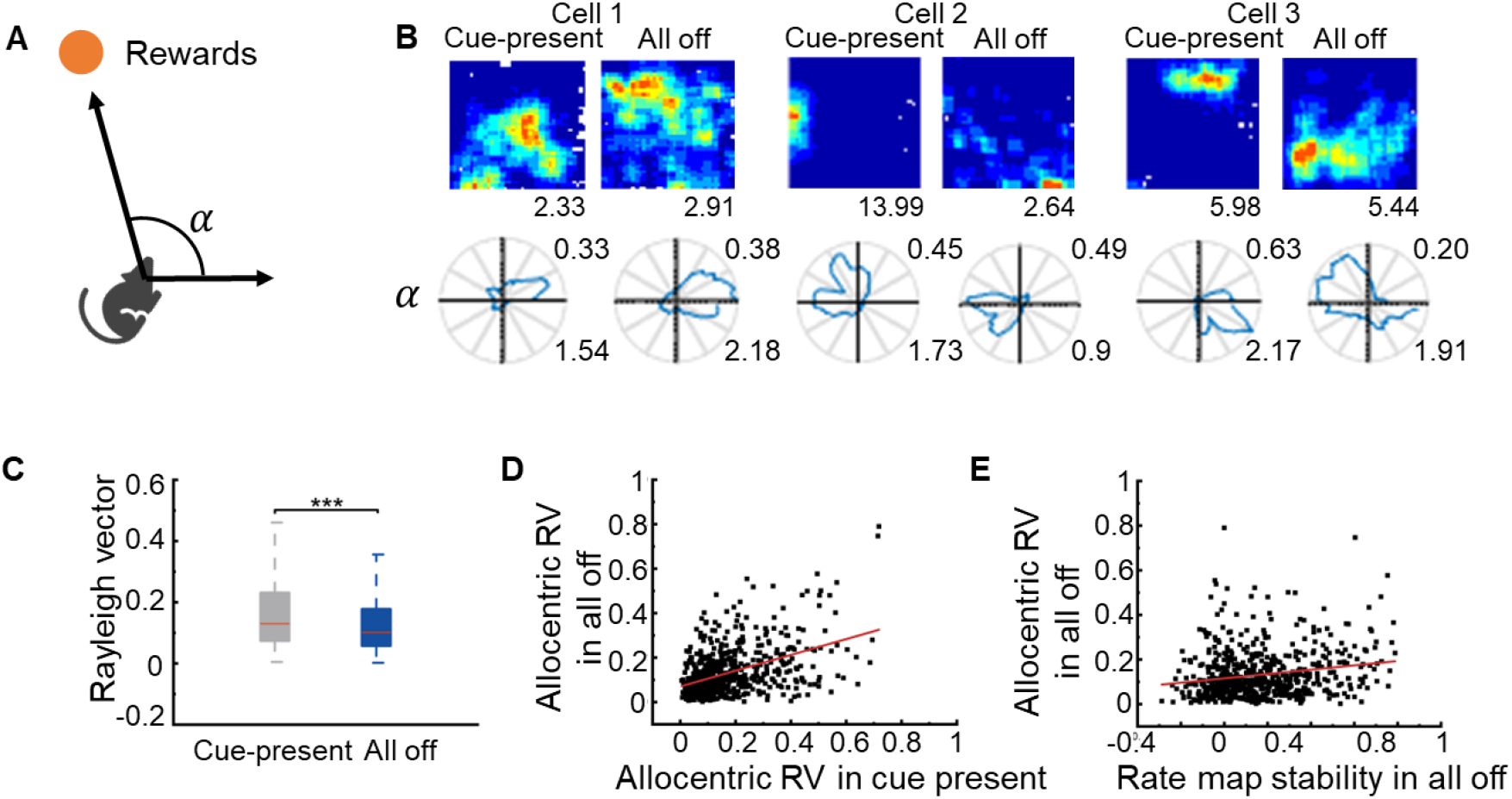
Place cell response to the locations of reward beacons. (A) Illustration of allocentric (*α*) direction relative to the location of a reward beacon. (B) Three example place cells show directional tuning towards the location of reward beacons. The upper row is rate maps and the lower row is allocentric reward-bearing polar plots. The numbers at the top right of polar plots are Rayleigh vector scores and the numbers at the bottom right are peak firing rates (Hz). (C) Rayleigh vector scores of allocentric tuning directions in the cue-present and All-Off conditions. Red lines are median, blue boxes cover the interquartile range *q*1*−q*3 and dashed lines cover *q*1 *−* 1.5 × IQR to *q*3 + 1.5 × IQR including all data points. (D) Relationship of allocentric Rayleigh vectors between the cue-present and All-Off conditions; (E) Relationship between the allocentric Rayleigh vectors and the intra-trial stability of rate maps in the All-Off condition. ^***^ p < 0.001.

## DISCUSSION

It has long been understood that the hippocampus uses multimodal sensory information to generate spatially-tuned neural firing and support spatial cognition^17–22^. Disentangling the contribution of different sensory modalities can be practically difficult in a real-world setting, hence the importance of introducing virtual reality techniques^5,9,23,24^, which allow arbitrary manipulations of sensory stimuli. In this study, we set out to investigate how purely visual stimuli can control and support place and grid cell firing, whilst mice are exploring a virtual reality environment^5,9^.

We have shown that purely visual barriers, which present no somatosensory or olfactory cues to the animal (unlike real-world barriers), nonetheless support place and grid cell firing. In the absence of these barrier cues (Wall-Off and All-Off manipulations), both place and grid cell firing is severely degraded. By contrast, a virtual floor alone cannot provide stability to spatially-tuned neurons. Our results are consistent with many lines of evidence demonstrating the importance of boundaries for neural representations of space: boundaries have a strong influence on shaping firing patterns of place cells and grid cells in rodents^6–8,25,26^, as well as neural representations in the human medial temporal lobes^27^.

Although the importance of boundaries to shape spatial firing is well recognised, no previous study has focused on which specific sensory properties of boundaries can modulate place and grid cell firing. The reported enhanced stability of grid cells in close proximity to boundaries^8^, for example, suggests that somatosensory information/physical contact with a barrier may be necessary to stabilise grid representations. Here we have shown, instead, that a purely visual boundary is sufficient for stabilising the firing of spatially-tuned neurons. Our study significantly adds to our understanding of the role of visual cues in supporting grid cell firing.

Grid cell networks display conserved internal dynamics across behavioural states^2,28,29^, and, when recorded in the absence of visual input, grid cells maintain consistent co-activity patterns, despite spatial instability^30^. Our results suggest that grid cell instability may be caused by excessive accumulation of path integration error, such that the internally coherent grid network ‘drifts’ with respect to the external world. This interpretation predicts that hexagonal grid patterns should be detectable at sufficiently short time scales, with the time scale depending on the rate of error accumulation. In contrast to the previous studies where only distance tuning, not drifting hexagonal patterns, could be detected when grid cell firing was disrupted in the dark^30,31^, our study is the first to demonstrate the presence of coherent but drifting hexagonal grid cell firing during spatial exploration, and shows that, in our experimental setup, error accumulation begins to degrade grid stability from 5 seconds onwards. Surprisingly, the presence of patterned visual cues on the floor of the virtual environment does not reduce the rate of error accumulation, indicating that not all visual cues can stabilise grid firing equally, and reinforcing the privileged role that boundaries play.

In addition to this, and unexpectedly, we observe that the firing patterns of place cells are more stable in areas near environmental boundaries even in the All-Off condition, when the boundaries are not directly visible, and the only environmental cue available is the (moving) reward beacon. We therefore hypothesise that the animal can infer the presence of the boundary from the movement of the beacon, relative to the animal’s own motor output. As the animal’s progress through the virtual space is blocked when it encounters the invisible boundary, this creates a mismatch between the visually observed beacon movement, and that predicted from the animal’s own motor output. We hypothesise that the animal can use this mismatch to infer boundary position, such that this inferred boundary can enhance place cell stability. To our knowledge, this is the first demonstration that an ‘invisible’ boundary, whose only physical manifestation is an impediment to movement, can modulate hippocampal spatial responses. Our results therefore expand the functional definition of boundaries to encompass entities that are not solely defined by their sensory characteristics.

Furthermore, our results show that the firing of a sub-population of place cells tracks the allocentric bearing of the animal to the reward beacon (which moves approximately every 4-5 seconds, but is stable between these relocations). This result is consistent with the previously reported modulation of hippocampal place cell firing by (stable) goals and/or objects^4,32–38^. We observe a significant correlation between place cell stability and the degree of their allocentric directional tuning to the reward beacon, indicating that tracking the allocentric location of the visible reward beacon enhances place cell stability.

In conclusion, our results demonstrate that purely visual boundaries can stabilise hippocampal spatial representations. We show, for the first time, the existence of coherently drifting hexagonal grid representations when boundary information is absent. Conversely, we also report the first evidence that hippocampal place cell responses can be modulated by sensorimotor feedback, on the basis of our unexpected observation that place cell responses can be stabilised by invisible boundaries.

## STAR METHODS

### Subjects and Surgery

Seven C57BL/6 mice were implanted with custom-made head plates and microdrives loaded with 17μm platinum-iridium tetrodes. Each mouse received a dual implant with one microdrive in right CA1 (ML: 1.8 mm, AP: 2.1 mm posterior to bregma) and the other in left mEC (ML = 3.1 mm, AP = 0.2 mm anterior to the transverse sinus, angled 4° posteriorly), each drive carrying four tetrodes. See further details in^5^. All procedures were carried out under the Animals (Scientific Procedures) Act 1986.

### Virtual Reality

Virtual environments (Figure S1) were constructed using a game engine (Unity, Unity Technologies) which ran on a Dell Precision T7500 workstation. The virtual scenes were rendered on a combination of four Acer B236HL LCD monitors mounted vertically in a square array plus two LCD projectors (native resolution 480 × 320, 150 lumens) mounted above. A mouse was attached to a head holder mounted in a bearing (Kaydon Reali-Slim bearing KA020XP0), which allowed free rotation on the horizontal plane. The bearing was held over the centre of an air-supported 200mm-diameter hollow polystyrene ball. The movement of the ball, triggered by the movement of the animal, can be detected by two optical computer mice (Logitech G700s gaming mouse) mounted with orthogonal orientations at the front and side of the ball. The output of the two optical mice drove the translational movement of the virtual scene in the X and Y axes respectively, corresponding to the movement of the animal. See further details in ^5,9^.

### Behavioural Training

After recovery from surgery, mice were also exposed to a 60cmx60cm square in a real environment to screen for place cells and grid cells. Electrodes were lowered by 62.5 μm per day independently on each side of the hemisphere until the spatial cells were found. In the meantime, mice were trained in the virtual environments as presented in^5,9^, learning to forage for the rewards indicated by a visual beacon. Foraging behaviour was motivated by sweetened soya milk drops as reward, delivered through a tube positioned within licking distance of the animal’s mouth. Visual beacons (a striped cylinder and a black rounded disk, see Figure S1) were placed in the virtual environments randomly to indicate the reward locations. At any given moment, there was always one reward location visually indicated by the beacon. The beacon would remain at the same location until mice visited the virtual rewarded location, at which point a drop of milk would be delivered. Once the reward was delivered, the beacon would be repositioned to a different random location, which could be at any position within the virtual environment, except within a radius of 20cm from the previous reward position.

A typical experimental session comprised two 40-min trials with a 40-min interval. Each trial consisted of either 40 min baseline trial or 20 min baseline and a 20 min manipulation trial. In the baseline trial - ‘Cue-present’, animals forage for rewards in a familiar 60cmx60cm virtual square environment identical to the one used during training. In the manipulation trial - ‘Cue-absent’, subsets of the visible cues in the virtual square environment were removed and replaced with a dark grey background (Figure S1C). The visual reward beacons were still available to indicate the location of reward, in all ‘Cue-absent’ trials.

### Data Acquisition

Extracellular action potentials were recorded using DACQ (Axona Ltd., UK). Spike sorting was performed offline using an automated clustering algorithm (KlustaKwik) followed by a manual review and editing step using an interactive graphical tool ‘Waveform’ (https://github.com/d1manson/waveform).

### Data Analysis

#### Classification of place cells and grid cells

Firing rate maps were constructed using 1.5 × 1.5 cm bins and a 5 × 5 boxcar filter. Spatial information and gridness scores were calculated using the methods ine^30^. Cells were classified as grid cells if their gridness scores in a cue-present condition exceeded the 95th percentile of a distribution of 1,000 gridness scores derived from spatially shuffled data, created by temporally shifting the spike train relative to position. Place cells were defined in a similar fashion, with spatial information being used to quantify the spatial tuning of a neuron.

#### Spike-triggered time-windowed rate maps

Time-windowed spatial displacement rate maps were constructed following the methods in ^39^. The gridness scores for the time-windowed maps were calculated by taking maximum gridness scores computed from rings as described as ‘Gridness measure 2’^40^. In brief, an autocorrelation of a smoothed rate map was first calculated and seven peaks closest to the centre were identified. The inner radius of the ring was defined as half of the mean distance from the peaks to the centre, and the outer radii ranged from the inner radius to the closest autocorrelogram edge. Then the rotational autocorrelations of these rings were calculated and a set of gridness scores were computed as the difference between the lowest correlation observed at 60 or 120 degrees of rotation and the highest correlation at 30, 90, or 150 degrees.

#### Place/Grid field stability

Intra-trial stability was calculated by correlating the firing rates of spatially corresponding bins between the first and the second halves of a trial. Inter-trial stability was measured by correlating the firing rates of spatially corresponding bins between the baseline and manipulation trials.

#### Analysis of allocentric reward-modulated firing

Allocentric reward-bearing polar plots were constructed by first calculating, for every position sample, the angle to the current reward position in the virtual environment reference frame. Angles were then binned into 6° bins, and spikes assigned to each bin. Both position maps and spike maps were smoothed separately using a 30° wide boxcar filter and finally the smoothed spike maps were divided by the smoothed position maps. The directional tuning of the cell was measured using the length of the mean resultant vector (Rayleigh Vector; RV) of the bins of the reward-tuned directional firing rate maps. To classify cells were as reward-tuned, the actual RV lengths were compared to null distributions of RV lengths derived from two different populations of spike-shuffled data, generated by: (1) shifting the entire spike train randomly with respect to position (as for place and grid cell classification), (2) maintaining the mean firing rate in each 2-dimensional rate map bin, but reassigning the spikes which occurred in each rate map bin randomly across the total set of time points for which the animal occupied that same bin. The second shuffle controlled for artifactual reward bearing tunings which may have arisen primarily from the concentration of firing in one particular (2-dimensional) location. A cell was classified as reward-tuned if its reward bearing map RV length exceeded the higher of the 99^th^ percentiles of the two shuffled reward bearing RV lengths described above.

#### Correlation between reward tuning and stability

To test whether the enhanced stability of reward tuned cells was primarily driven by their proximity to a boundary, the correlation of stability versus reward tuning was a partial correlation, controlling for a ‘boundary score’, which quantified the extent to which firing occurred near a boundary. The boundary score was calculated by assigning weights to each bin in a rate map based on its distance to the closest edge. The boundary score for each cell was defined by the sum of the firing rates within each bin multiplied by the corresponding weight from the mask.

## ACKNOWLEDGMENTS

This study was funded by the BBSRC (BB/W007878/1 to G.C. and BB/I021221/1 to F.C.), the Royal Society (RGS\R1\221070 to G.C.), the European Union H2020 research and innovation programme (Human Brain Project #785907, SGA2 to G.C., F.C., N.B.), China Scholarship Council (X.Y., G.C.), the Wellcome Trust (Fellowship 220886/Z/20/Z to T.W.), the European Research Council (consolidator award ‘DEVMEM’ to F.C.) and the Wellcome Trust (Investigator Award, 210690/Z/18/Z).

## AUTHOR CONTRIBUTIONS

G.C. and F.C. conceived the study. N.B. and G.C. constructed and implemented the virtual reality setup. G.C. conducted the experiments and spike-sorted the data. X.Y., T.W. and G.C. analyzed the data. T.W., F.C. and G.C. wrote the manuscript with input from N.B.

## SUPPLEMENTAL INFORMATION

**Figure S1.**
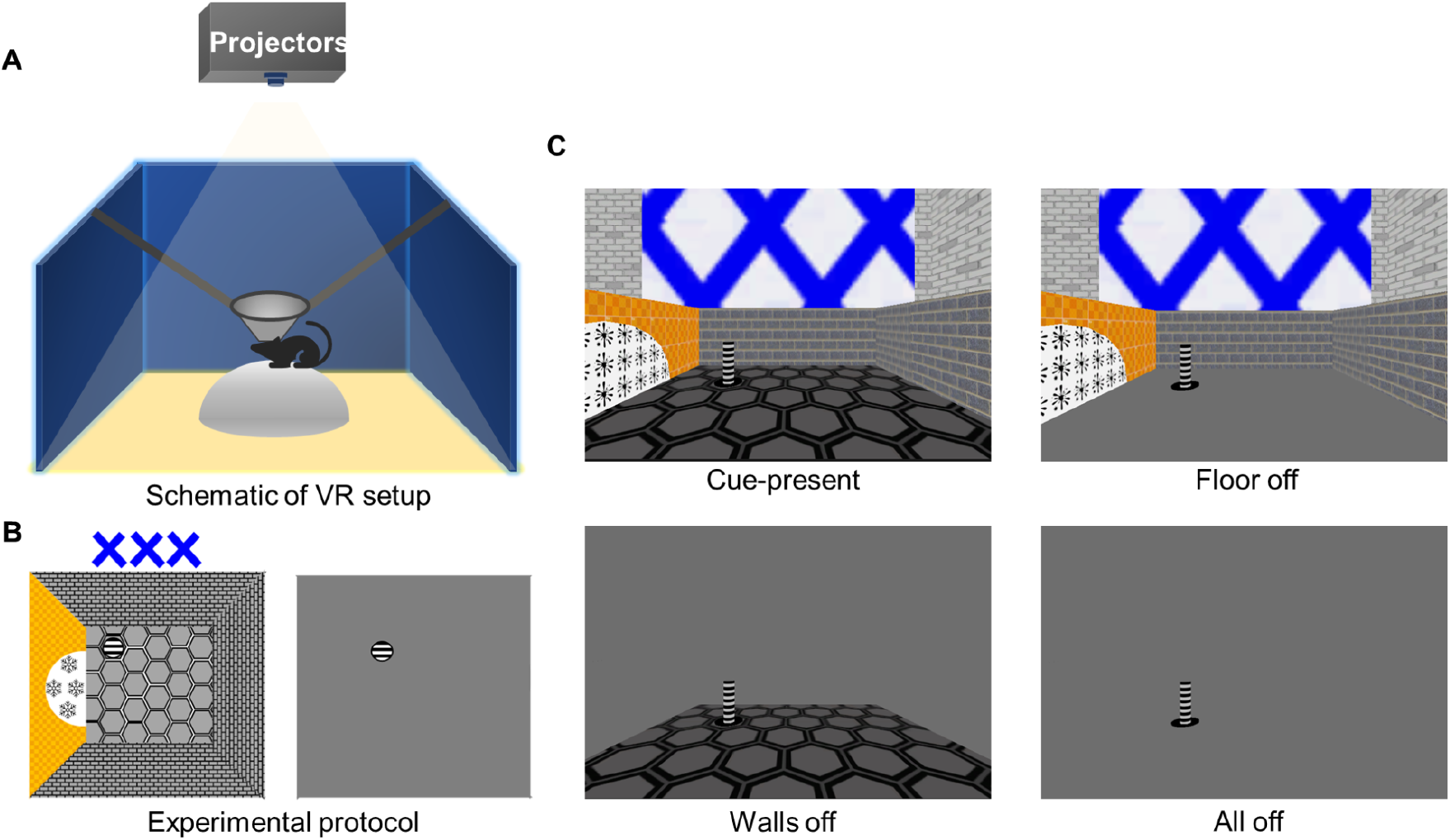
VR setup and experimental protocol. (A) Schematic of the virtual reality setup. (B) Experimental protocol: a cue-present trial with all visual cues being visible (left) and a cue-absent (All Off) trial with all visual cues being invisible apart from the visual beacon (right). (C) Side views of the VR environment in the baseline/cue-present condition (top left) and all three cue-absent conditions: Floor Off (top right), Walls Off (bottom left), All Off (bottom right).

**Figure S2.**
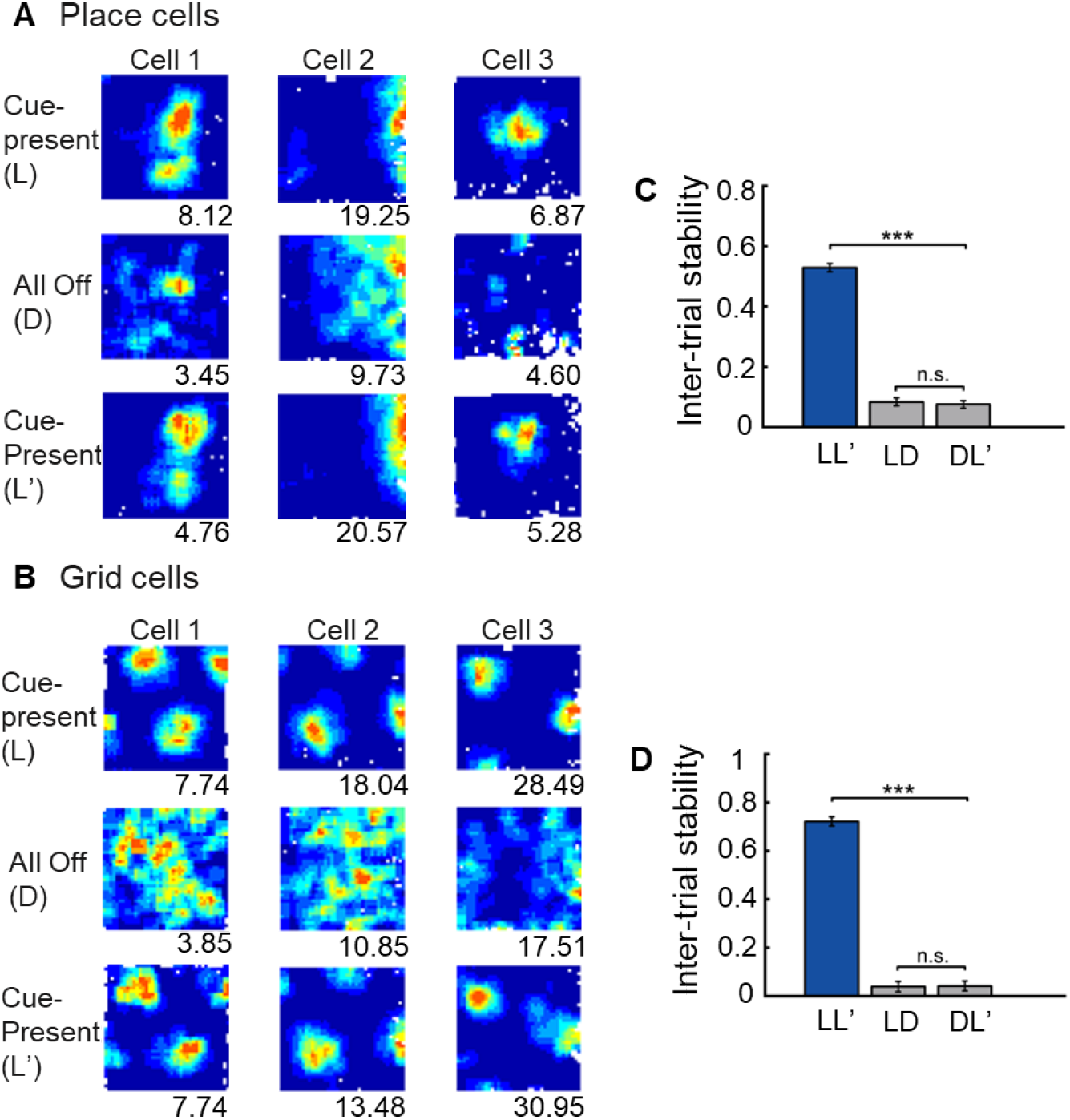
Stable recordings of place cells and grid cells. (A) Rate maps for three representative place cells recorded in a 60 cm virtual square. (B) Rate maps for three representative grid cells in the same virtual square. Numbers at the bottom right of rate maps are peak firing rate (Hz). The top rows are from the 1st baseline trials in the cue-present (Light, L) condition, the second rows are the trials in the All-Off (Dark, D) condition and the last rows are the 2nd baseline trials. (C) Inter-trial stability of place cells between the first and the second cue-present trials (LL’), the first cue-present trial and the All-Off trial (LD), and the All-Off trial and the second cue-present trial (DL’). The inter-trial stability between the two baseline trials (LL’) is significantly higher than that between the cue-present and the All-off trials: LD and DL’ (Paired t-test, t(322) = 25.09, p<0.001; t(322) = 27.19, p < 0.001); there is no significant difference between LD and DL’ (Paired t-test, t(322) = 0.91, p = 0.36). (D) Inter-trial stability of grid cells between the first and the second cue-present trial (LL’), the first cue-present trial and the All-Off trial (LD), and the All-Off trial and the second cue-present trial (DL’). The inter-trial stability between the two baseline trials is significantly higher than that between LD and DL’ (Paired t-test, t(58) = 26.82, p < 0.001; t(58) = 26.63, p < 0.001); there is no significant difference between LD and DL’ (Paired t-test, t(58) = -0.2, p = 0.84).

**Figure S3.**
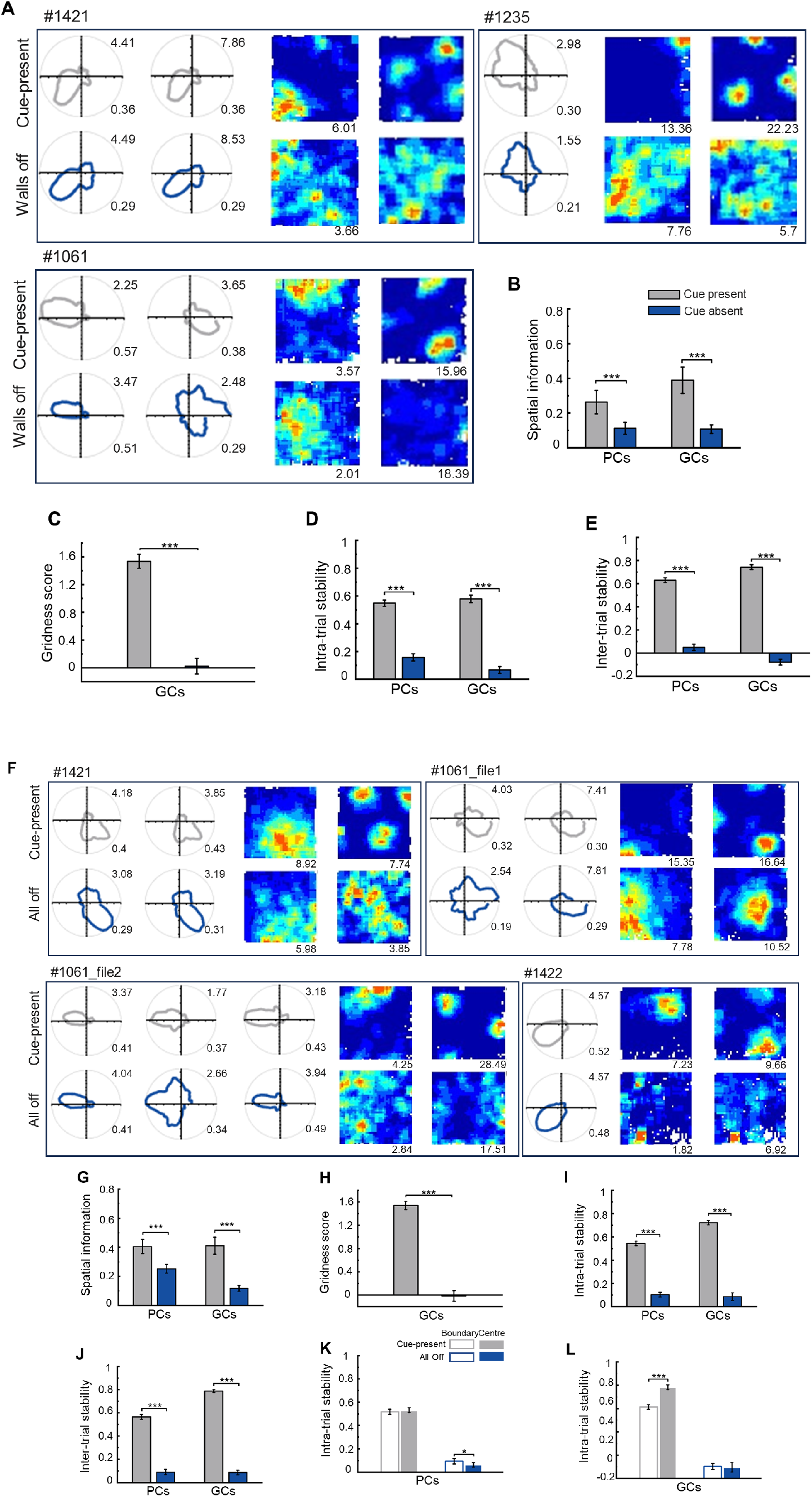
Analysis of spatial firing for place and grid cells co-recorded with mEC Head Direction (HD) cells. Only HD cells with RV>0.3 in the cue-present trials were included. Head direction cells maintained their directional tuning in Walls-Off (A) and All-Off conditions (F). Each panel shows example plots for a selection of co-recorded cells: the polar plots on the left show responses of co-recorded head direction cells; the ratemaps on the right spatial maps for one co-recorded place cell and one co-recorded grid cell. For polar plots, numbers at the top right are Rayleigh vector scores; numbers at the bottom right are peak firing rates (Hz). Numbers on top of the panels are animal numbers. (B-E) Comparison of firing properties of place cells and grid cells co-recoded with HD cells including spatial information (B, paired t-test: place cells, t(64) = 4.07, p < 0.001; grid cells, t(27) = 5.74, p < 0.001), gridness scores (C, paired t-test: t(27) = 5.61, p < 0.001), intra-trial stability (D, paired t-test: place cells, t(64) = 10.15, p < 0.001; grid cells, t(27) = 16.52, p < 0.001) and inter-trial stability (E, paired t-test: place cells, t(64) = 17.35, p < 0.001; grid cells, t(27) = 23.38, p < 0.001) between the cue-present and Walls-Off trials. (G-L) Comparison of firing properties of place cells and grid cells co-recoded with HD cells including spatial information (G, paired t-test: place cells, t(123) = 8.71, p < 0.001; grid cells, t(45) = 7.84, p < 0.001), gridness scores (H, paired t-test: t(45)=10.50, p<0.001), intra-trial stability (I, paired t-test: place cells, t(123) = 16.84, p < 0.001; grid cells, t(45) = 14.34, p < 0.001), inter-trial stability (J, paired t-test: place cells, t(123) = 14.53, p < 0.001; grid cells, t(45) = 29.84, p < 0.001), intra-trial stability of boundary (open bars) and central (solid bars) rate maps for place cells (K, ANOVA Zone*Cue presence, F(1,123) = 0.05, p = 0.829; paired t-test: All-Off, t(123) = 1.87, p < 0.05; cue-present, t(123) = 1.44, p = 0.076), Intra-trial stability of boundary (open bars) and central (solid bars) rate maps in cue-present (grey) for grid cells (L, ANOVA Zone*Cue presence, F(1,45) = 11.52, p = 0.001; paired t-test: All-Off, t(45) = -0.14, p = 0.893; cue-present, t(45)= -6.57, p < 0.001) between the cue-present and All-Off trials. ^*^p < 0.05; ^**^ p < 0.01; ^***^ p < 0.001.

